# More accurate analysis of maternal effect genes by siRNA electroporation into mouse oocytes

**DOI:** 10.1101/2022.10.31.514463

**Authors:** Takuto Yamamoto, Shinnosuke Honda, Issei Ideguchi, Motoki Suematsu, Shuntaro Ikeda, Naojiro Minami

## Abstract

Maternal RNAs and proteins accumulate in mouse oocytes and control the initial stages of development. The DNA in sperm combines with protamine, which is exchanged after fertilization with maternal histones, including H3.3, but the effect of H3.3 on post-fertilization development has been unclear. In this study, we established an electroporation method to introduce H3.3 siRNA into germinal vesicle (GV)-stage oocytes without removing cumulus cells. In the traditional microinjection method, cumulus cells attached to oocytes must be removed; however, we confirmed that artificially removing cumulus cells from oocytes reduced fertilization rates, and oocytes originally free of cumulus cells had reduced developmental competence. When H3.3 siRNA was introduced at the GV stage, H3.3 was still present in the maternal pronucleus and second polar body, but not in the paternal pronucleus, resulting in embryonic lethality after fertilization. This indicates that the H3.3 protein was not incorporated into the paternal pronucleus because it was repeatedly translated and degraded in a relatively short time. On the other hand, H3.3 protein incorporated into the maternal genome in the GV stage escaped degradation and remained in the maternal pronucleus after fertilization. This new method of electroporation into GV-stage oocytes without removing cumulus cells is not skill intensive and is essential for the accurate analysis of maternal effect genes.

**In brief:** For analysis of maternal factors contained in the oocyte, suppression of genes within the oocyte is necessary. In this paper, siRNA was introduced into oocytes by electroporation, showing that maternal mRNA was suppressed efficiently.

## Introduction

In mammals, maternal transcripts and proteins (maternal factors) that support early embryonic development accumulate during oogenesis. Since there is little transcriptional activity during the period from the development of fully grown oocytes to that of early one-cell embryos in mice (Bachvarova, 1985; Aoki *et al*., 1997; De La Fuente and Eppig, 2001), these transcripts and proteins are considered to regulate the developmental program during and after this period. Some of these factors have already been shown to be involved in meiotic progression, maternal RNA degradation, and chromatin remodeling (Minami et al., 2007; Miyamoto, 2013; Daniel E Shumer, 2017). Maternal transcripts are generally degraded around the two-to eight-cell stage in mice, which coincides with zygotic genome activation (Tadros and Lipshitz, 2009).

Knockout and transgenic mice are often used to analyze the function of maternal factors (Tong et al., 2000; Wu et al., 2003; Rajkovic et al., 2004; Honda et al., 2018). However, these genetically modified techniques are very time consuming, and they require the management of large numbers of animals. Subsequently, a method of microinjecting siRNA directly into germinal vesicle (GV)-stage oocytes or metaphase II (MII) oocytes has been developed, making it possible to analyze maternal factors in just one generation (Inoue et al., 2012; Inoue and Zhang, 2014). Although cumulus cells attached to oocytes must be removed by pipetting prior to microinjection, it is unclear how the oocytes are affected by this procedure. Electroporation was developed as an alternative gene delivery method to microinjection, the latter of which requires skilled micromanipulation techniques (Peng et al., 2012a). Electroporation is often used as a tool to introduce Cas9-gRNA complexes, morpholino oligonucleotides, and plasmid DNAs into zygotes (Peng et al., 2012a; Kaneko et al., 2014). Although a method has also been established to introduce siRNA into GV-stage oocytes by electroporation, this technique is very complicated because it requires removing the cumulus cells, thinning the zona pellucida, and using polyamine-based transfection reagents (Peng et al., 2012b). It is known that two types of GV-stage oocytes can be artificially harvested from mouse ovaries: those surrounded by cumulus cells (cumulus-enclosed oocytes: CEOs) and those without associated cumulus cells (denuded oocytes: DOs) (De La Fuente and Eppig, 2001). DOs have a lower maturation rate and higher transcriptional activity than CEOs (Liu and Aoki, 2002; Kikuchi *et al*., 2016), but the differences in subsequent fertilization and development rates between DOs, CEOs, and artificially denuded CEOs (dCEOs) have not been well studied. Low numbers of cumulus cells surrounding oocytes have been associated with follicle atresia in humans, cynomolgus monkeys, cattle, and horses (Gougeon and Testart, 1986; Lefevre et al., 1988; Blondin and Sirard, 1995; Hinrichs and Williams, 1997), which may be a reason for the poor quality of DOs.

In mammals, there are three non-centromeric histone H3 variants (H3.1, H3.2, and H3.3), and H3.3 is known to be a maternal factor. H3.3 is incorporated into the paternal pronucleus immediately after fertilization, and also promotes chromatin decondensation (Torres-Padilla et al., 2006; Akiyama et al., 2011; Lin et al., 2013, 2014; Inoue and Zhang, 2014)·H3.3 accumulates in active chromatin after the two-cell stage in mouse embryos and is thought to regulate embryo-specific gene expression patterns (Ishiuchi et al., 2021; Aoki, 2022). Although several H3.3 knockdown experiments in MII and GV-stage oocytes have been reported (Inoue and Zhang, 2014; Lin et al., 2014; Wen et al., 2014a; Kong et al., 2018), the post-fertilization developmental competence of H3.3-depleted GV-stage oocytes has not been examined.

In this study, we investigated in detail the differences in fertilization and development rates between DOs, CEOs, and dCEOs. CEOs had higher fertility and developmental competence than DOs. We also found that cumulus cells were essential for CEO fertility, which led us to develop a method that used electroporation to introduce maternal transcript-targeting siRNAs into oocytes without removing the cumulus cells. This electroporation method allowed us to investigate H3.3 function more precisely in embryonic development after fertilization.

## Methods

### Collection of GV-stage oocytes

GV-stage oocytes were obtained from 8- to 12-week-old ICR mice (Japan SLC) 48 h after injection with 7.5 I.U. eCG. The ovaries were removed and transferred to M2 medium containing 0.2 mM 3-isobutyl-1-methylxanthine (IBMX; Nacalai Tesque). The ovarian follicles were punctured with a 26-gauge needle, and oocytes with homogenous cytoplasm were collected. Collected oocytes were grouped as follows: those with more than three layers of unexpanded cumulus cells (CEOs), those in which approximately half of the cumulus layers were intact (partially denuded oocytes: PDOs), and those without cumulus cells (DOs). dCEOs and denuded PDOs (dPDOs) were prepared by mechanically removing cumulus cells from CEOs and PDOs with a narrow pipette, and pseudo-denuded DOs were subjected to the same manipulation. CEOs, PDOs, and DOs are shown in Fig. 1.

**Fig. 1.**
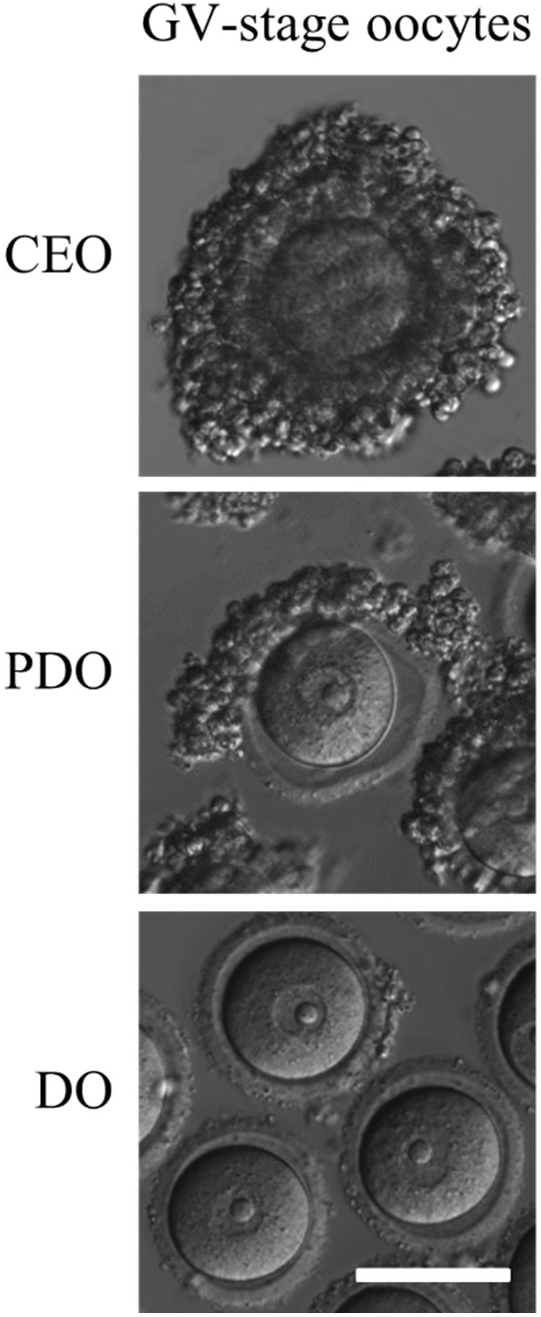
Morphology of germinal vesicle (GV)-stage oocytes collected from mouse ovaries. CEO: cumulus-enclosed oocytes. PDO: partially denuded oocytes. DO: denuded oocytes. Scale bar, 100 μm.

### Introduction of siRNA into GV-stage oocytes by electroporation

Electroporation of oocytes was performed with the NEPA21 electroporator system (NEPA GENE) using a glass slide on which there were two metal electrodes separated by a 1-mm gap (CUY501P1-1.5; NEPA GENE). Electroporation parameters consisted of four poring pulses (40 V; pulse length, 1.0, 2.0, 3.0, 4.0, or 8.0 ms; interval, 50 ms; decay rate, 10%; polarity, +) and five transfer pulses (5 V; pulse length, 50 ms; interval, 50 ms; decay rate, 40%; polarity, +/-). Opti-MEM (Thermo Fisher Scientific) containing siH3.3 (4 siRNAs at 5 μM each; RNAi Inc.) or 4 μM red fluorescent non-targeting siRNA (BLOCK-iT™ Alexa Fluor^®^ Red Fluorescent Control; Thermo Fisher Scientific) was prepared as siRNA solution. The siRNA solution (5 μl) was applied between the two electrodes on the glass slide. Oocytes were placed in a line between the electrodes, and the electrical discharge was performed. The siRNA solution was exchanged every two operations to avoid dilution of the solution. After electroporation, the oocytes were transferred into α-MEM supplemented with 5% fetal bovine serum and 10 ng/ml epidermal growth factor to induce meiotic maturation. In this experiment, *H3f3a* and *H3f3b* were targeted using the following siRNA sequences, as in a previous report (Wen *et al*., 2014b): *siH3f3a* 1s, CGUUCAUUUGUGUGUGAAUUUtt; siH3f3a 1as, AAAUUCACACACAAAUGAACGtt; siH3f3a 2s, GCGAGAAAUUGCUCAGGACUUtt; siH3f3a 2as, AAGUCCUGAGCAAUUUCUCGCtt; *siH3f3b* 1s, UCUGAGAGAGAUCCGUCGUUAtt; siH3f3b 1as, UAACGACGGAUCUCUCUCAGAtt; siH3f3b 2s, GAAGCUGCCAUUCCAGAGAUUtt; and siH3f3b 2as, AAUCUCUGGAAUGGCAGCUUCtt. The following were also used: control siRNA s, UACGAAUGACGUGCGGUACGU; and control siRNA as, GUACCGCACGUCAUUCGUAUC.

### In vitro fertilization and embryo culture

After *in vitro* maturation for 15–18 h, MII oocytes were transferred into a 100-μL droplet of human tubal fluid (HTF) medium supplemented with 4 mg/mL bovine serum albumin (BSA; Sigma-Aldrich) (Minami *et al*., 2001). Spermatozoa were collected from the cauda epididymis of 12- to 18- week-old ICR male mice. After 1-h preincubation in HTF medium, sperm suspension was added into fertilization droplets at a final concentration of 1×10^6^ cells/mL. Six hours post-insemination (hpi), fertilized embryos were washed in potassium simplex optimized medium (KSOM) supplemented with amino acids (Ho *et al*., 1995) and 1 mg/mL BSA, and then cultured in the same medium under paraffin oil (Nacalai Tesque). All incubations were performed at 37°C under 5% CO_2_ in air.

### RNA extraction and reverse transcription-quantitative PCR (RT-qPCR)

Total RNA extraction and the synthesis of cDNA from oocytes or cumulus cells were performed using the superPrep™ Cell Lysis & RT Kit for qPCR (TOYOBO). Synthesized cDNA was mixed with specific primers and KOD SYBR qPCR Mix (TOYOBO), then amplified by RT-qPCR. The protocols for RT-qPCR and determination of transcript levels were previously described (Shikata et al., 2020), and Gapdh was used as an internal control. Relative gene expression was calculated by the 2^-ΔΔCt^ method (Livak and Schmittgen, 2001). The primer sequences used for RT-qPCR were as follows: Gapdh, 5’-GTGTTCCTACCCCCAATGTG-3’ (forward) and 5 TGTCATCATACTTGGCAGGTTTC-3’ (reverse); H3f3a, 5’-ACAAAAGCCGCTCGCAAGAG-3’ (forward) and 5’-ATTTCTCGCACCAGACGCTG-3’ (reverse); and H3f3h, 5’- TGGCTCTGAGAGAGATCCGTCGTT-3’ (forward) and 5’-GGATGTCTTTGGGCATGATGGTGAC-3’ (reverse).

### Immunocytochemistry and fluorescence analysis

To detect nuclear localization of H3.3, oocytes and embryos were fixed with 3.2% paraformaldehyde and 0.2% Triton X-100 (Sigma-Aldrich) in phosphate-buffered saline (PBS) for 20 min at 28°C. Oocytes and embryos were blocked in poly(butylene succinate-co-terephthalate) (PBST) containing 1.5% BSA, 0.2% sodium azide, and 0.02% Tween20 (antibody dilution buffer) for 1 h at 28°C, then incubated overnight at 4°C with rat anti-H3.3 (1:100; CE-040B; Cosmo Bio) antibody in antibody dilution buffer. The samples were then washed in PBST and incubated with Alexa Fluor 555 donkey anti-rat IgG secondary antibody (1:500; Thermo Fisher Scientific) for 1 h at 28°C. After washing in PBST, nuclei were stained in PBST containing 10 μg/mL Hoechst 33342 (Sigma-Aldrich) for 10 min at 28°C. Stained samples were mounted on slide glasses, and signals were observed using a fluorescence microscope (IX73; Olympus). Similarly, GV-stage oocytes were mounted on slide glasses, and fluorescence signals of BLOCK-iT™ Alexa Fluor^®^ Red Fluorescent Control (Thermo Fisher Scientific) were then observed using a fluorescence microscope (BX50; Olympus).

### Statistical analyses

Developmental rates were analyzed by the chi-square test, with Holm’s adjustment. In gene expression analysis, RT-qPCR data were analyzed by Student’s t-test for pairwise comparisons or by one-way analysis of variance (ANOVA) followed by Tukey-Kramer test for multiple comparisons. P values <0.05 were considered to be statistically significant.

### Ethical approval for the use of animals

All experimental procedures were approved by the Animal Research Committee of Kyoto University (Permit no. R3-17) and were performed in accordance with the committee’s guidelines.

## Results

The existence of cumulus cells affects the developmental competence of mouse GV-stage oocytes The method used to introduce siRNA into mouse GV-stage oocytes was first investigated by determining how the developmental competence of each GV oocyte differed depending on cumulus cell status. Three types of GV-stage oocytes were harvested from mouse ovaries: CEOs, PDOs, and DOs. In the first experiment, in vitro maturation (IVM) and subsequent in vitro fertilization (IVF) were performed on these three cell types to determine differences in their fertilization and developmental rates. At 6 hpi, embryos with two pronuclei were defined as fertilized embryos. Although there was no difference between their fertilization rates, development to the blastocyst stage occurred at a significantly lower rate in DOs than in CEOs (Table 1), which is consistent with previous reports (Shioya et al., 1988; Liu and Aoki, 2002; Kikuchi et al., 2016). In the next experiment, the differences in the fertilization and developmental rates of CEOs, PDOs, and DOs were compared to those of dCEOs and dPDOs. The denudation procedure was performed using a pipette. Pseudo-denuded DOs were checked to confirm the effects of the pipetting procedure on cumulus cell denudation, and there was no difference in the fertilization or developmental rate between DOs and pseudo-denuded DOs (data not shown). To further investigate the effects of denudation and the absence of cumulus cells on fertilization and subsequent development, the removed cumulus cells were co-cultured with dCEOs and DOs during IVM and IVF. The fertilization rates of dCEOs and dPDOs were dramatically reduced, but the rate partially recovered when dCEOs were co-cultured with cumulus cells and developed into blastocysts (Table 1). Co-culture with cumulus cells did not affect the fertilization or developmental rate of DOs.

**Table 1.** Effects of attached cumulus cells on developmental competence of CEOs, PDOs, and DO.

| Oocyte | No. of<br>GV-stage oocytes | No. (%) of embryos<br>with two pronuclei | No. (%) of<br>2-cell embryos | No. (%) of<br>blastocysts | % of blastocysts<br>per fertilized embryo |
| --- | --- | --- | --- | --- | --- |
| CEO | 103 | 67 (65.0)<sup>a</sup> | 66 (64.1)<sup>a</sup> | 56 (54.4)<sup>a</sup> | 83.6<sup>a</sup> |
| dCEO | 102 | 3 (2.94)<sup>c</sup> | 3 (2.94)<sup>d</sup> | 0 (0.0)<sup>d</sup> | 0.0<sup>ab</sup> |
| dCEO + cumulus cells | 59 | 12 (20.3)<sup>b</sup> | 13 (22.0)<sup>c</sup> | 10 (16.9)<sup>c</sup> | 83.3<sup>ab</sup> |
| PDO | 98 | 70 (71.4)<sup>a</sup> | 68 (69.4)<sup>a</sup> | 42 (42.9)<sup>ab</sup> | 60.0<sup>ab</sup> |
| dPDO | 117 | 44 (37.6)<sup>b</sup> | 38 (32.5)<sup>bc</sup> | 19 (16.2)<sup>c</sup> | 43.2<sup>b</sup> |
| DO | 117 | 69 (59.0)<sup>a</sup> | 58 (49.6)<sup>ab</sup> | 36 (30.8)<sup>bc</sup> | 52.2<sup>b</sup> |
| DO + cumulus cells | 104 | 70 (67.3)<sup>a</sup> | 62 (59.6)<sup>a</sup> | 49 (47.1)<sup>ab</sup> | 70.0<sup>ab</sup> |

Different letters in the same column (a, b, c, and d) represent significant differences ( $P$ value < 0.05).

Assessing the electroporation of GV-stage oocytes using red fluorescent non-targeting siRNA Electroporation was performed on CEOs and DOs as a method for introducing siRNA into GV-stage oocytes without removing the cumulus cells. To visualize the incorporation of siRNA into GV-stage oocytes, a red fluorescent non-targeting siRNA (BLOCK-iT^™^ Alexa Fluor^®^ Red Fluorescent Control; Thermo Fisher Scientific) was introduced into CEOs and DOs by electroporation under various pulse conditions (pulse length: 0, 1.0, 2.0, 3.0, and 4.0 ms). Efficient incorporation of siRNA into GV-stage oocytes was confirmed at a pulse length of 4.0 ms for both CEOs and DOs (Fig. 2).

**Fig. 2.**
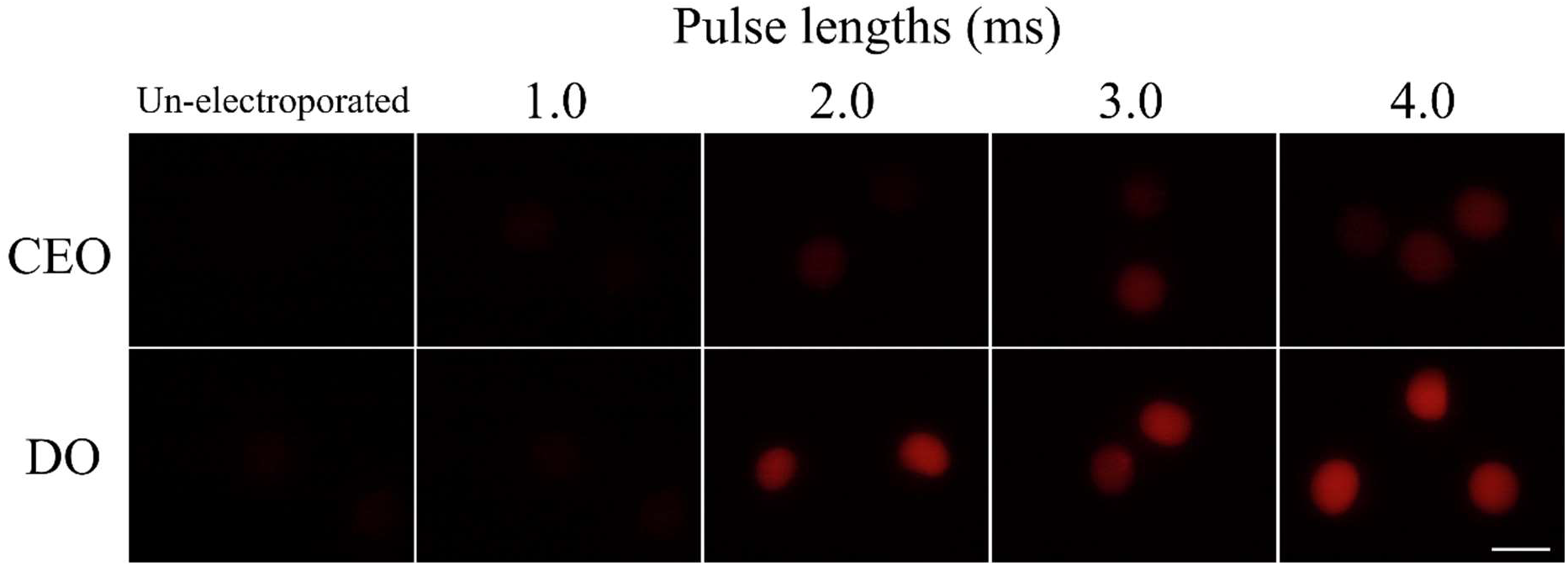
Effects of pulse length on siRNA incorporation into cumulus-enclosed oocytes (CEOs) and denuded oocytes (DOs). Representative images of germinal vesicle (GV)-stage oocytes electroporated with red fluorescent non-targeting siRNA under various pulse lengths are shown. Scale bar, 100 μm.

### The electroporation of siH3.3 effectively reduces the mRNA level of MII oocytes

To investigate the efficacy of the electroporation method for suppressing specific genes, siRNAs targeting *H3f3a* and *H3f3b*, which encode histone H3.3, were introduced into GV-stage oocytes by electroporation (two siRNAs per gene, a total of four) at pulse lengths of 1.0, 4.0, and 8.0 ms. Immediately after electroporation, IVM was performed for 15 h, and the efficacy of suppressing gene expression was examined by RT-qPCR using oocytes that reached the MII stage. The expression of both *H3f3a* and *H3f3b* was suppressed by more than 70% compared to control cells electroporated with siControl, a non–gene targeting siRNA (Fig. 3A). The 1.0-, 4.0-, and 8.0-ms pulse lengths resulted in comparable knockdown efficacy. Gene expression was also suppressed in cumulus cells attached to CEOs, suggesting that siRNA was introduced into cumulus cells (Fig. 3B).

**Fig. 3.**
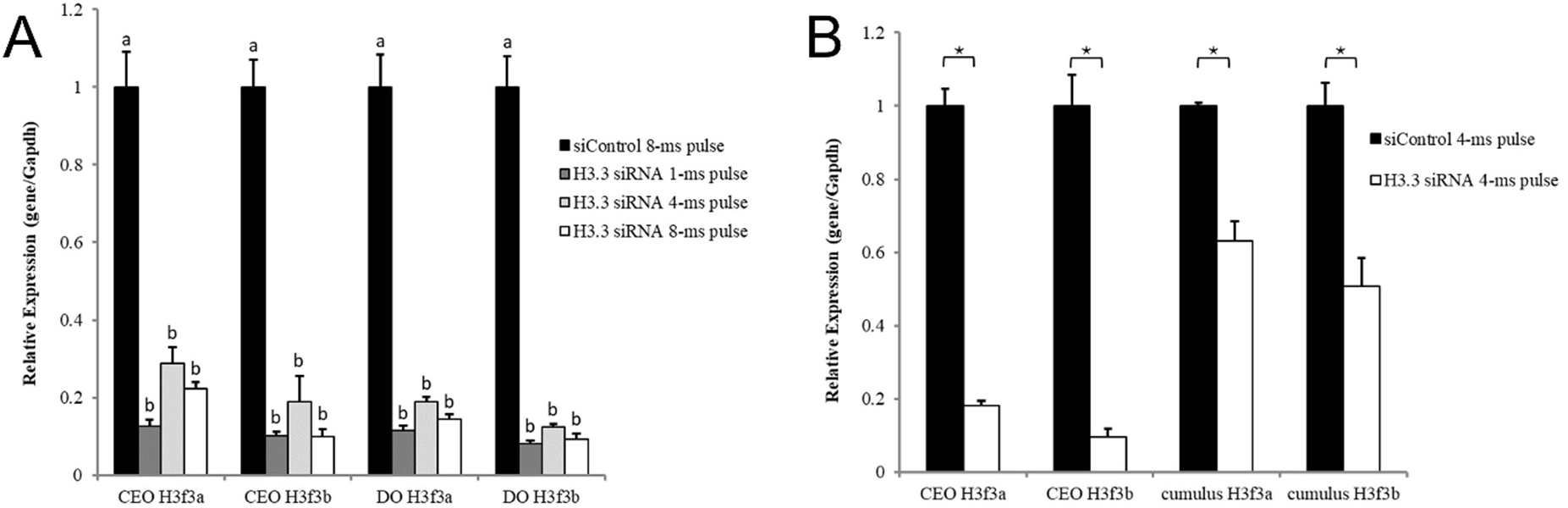
Reverse transcription-quantitative PCR (RT-qPCR) analysis of H3f3a and H3f3b mRNA in H3.3-knockdown oocytes at the metaphase II (MII) stage. (A) RT-qPCR of H3f3a and H3f3b was performed using electroporated cumulus-enclosed oocytes (CEOs) and denuded oocytes (DOs) after in vitro maturation (IVM). Gene expression levels were normalized to Gapdh as an internal control. Data are expressed as means ± S.E.M (n=3). Fifteen embryos were analyzed. Statistical analysis was performed using one-way analysis of variance (ANOVA) followed by Tukey-Kramer test. P values < 0.05 were considered to be statistically significant and are represented by different letters (a and b). (B) RT-qPCR of H3f3a and H3f3b was performed using cumulus cells attached to electroporated CEOs after IVM. Gene expression levels were normalized to Gapdh as an internal control. Data are expressed as means ± S.E.M (n=3). Statistical analysis was performed using Student’s t-test. P values < 0.05 were considered to be statistically significant.

### H3.3 is incorporated into the nucleus of GV-stage oocytes and its incorporation into the paternal pronucleus can be blocked after fertilization

To monitor the expression of H3.3 before and after fertilization and to confirm the reduction of H3.3 protein by siRNA treatment, we performed immunofluorescence staining for H3.3 from the GV stage to the one-cell stage. H3.3 was already incorporated into the nuclei of GV-stage oocytes (Fig. 4A). In MII oocytes, H3.3 was incorporated into both control and H3.3 knockdown oocytes (Fig. 4B). At 6 hpi, H3.3 was incorporated into maternal and paternal pronuclei and the nuclei of the two polar bodies in control embryos. On the other hand, H3.3 was not incorporated into paternal pronuclei of H3.3 knockdown embryos (Fig. 4B).

**Fig. 4.**
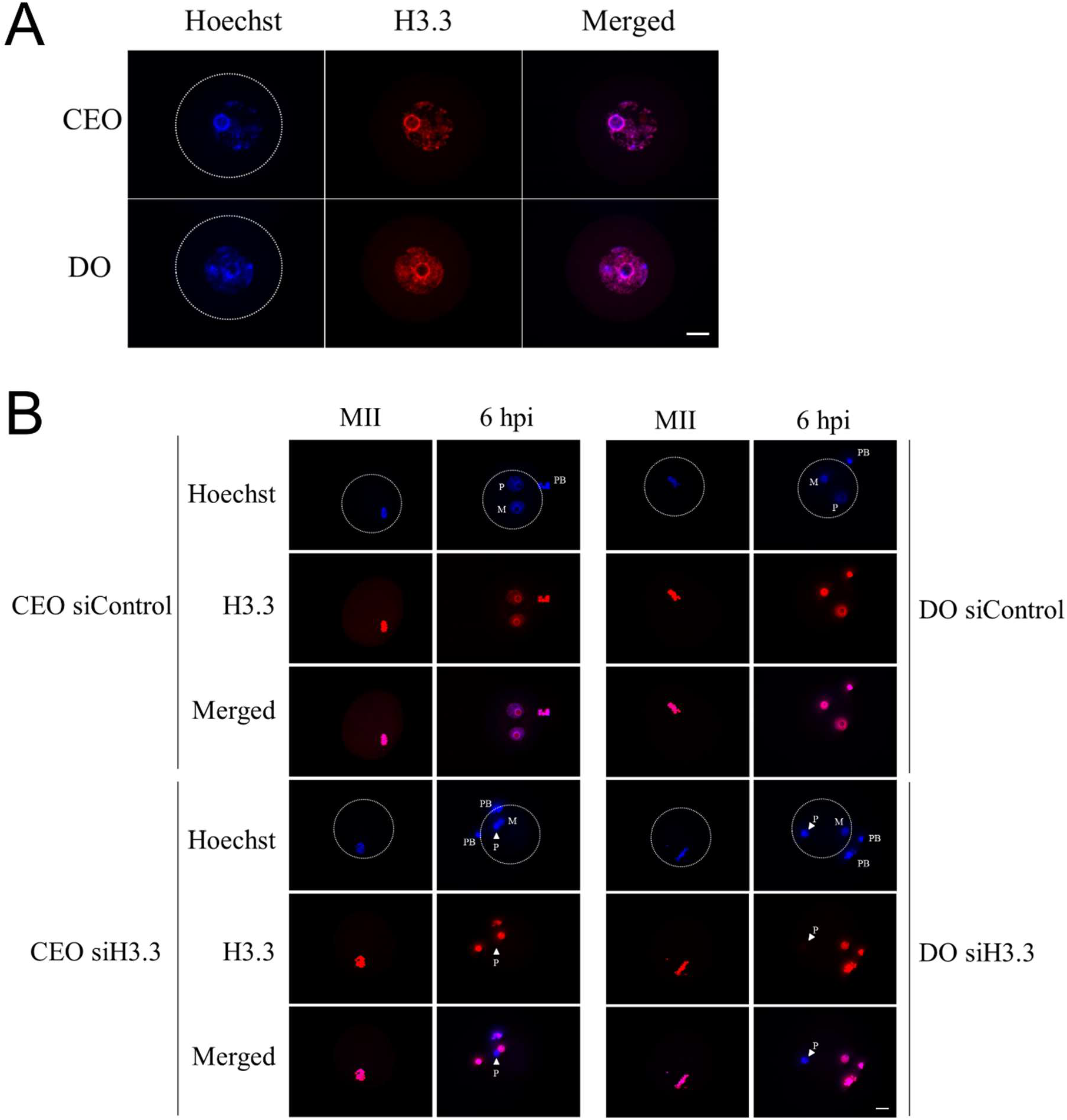
Effects of siH3.3 treatment on H3.3 incorporation into germinal vesicle (GV)-stage oocytes before and after fertilization. Oocytes and embryos were stained with anti-H3.3 antibody. (A) Localization of H3.3 in cumulus-enclosed oocytes (CEOs) and denuded oocytes (DOs). Cumulus cells of CEOs were removed for immunostaining. H3.3 was present in the nuclei of GV-stage oocytes and cumulus cells. Scale bar, 20μm. (B) Localization of H3.3 in metaphase II (MII) oocytes and one-cell embryos. H3.3 is localized in the nuclei of MII oocytes treated with or without siH3.3. However, H3.3 is not localized in the paternal pronuclei of embryos at 6 hpi produced after treatment with siH3.3 at the GV stage. Scale bar, 20 μm.

### The developmental competence was reduced by electroporation of siH3.3 into GV-stage oocytes

H3.3 is present in the oocyte as a maternal factor, and it is incorporated into the paternal pronucleus immediately after fertilization. However, it is unclear how removing maternal H3.3 from GV-stage oocytes affects post-fertilization development (van der Heijden et al., 2005; Torres-Padilla et al., 2006; Akiyama et al., 2011). CEOs and DOs electroporated with siH3.3 or siControl with a 1.0-ms pulse length were subjected to IVM for 15 h to examine maturation rates and the differences in fertilization and developmental rates after IVF (Table 2). siH3.3 did not affect the maturation or fertilization rate, but the rate of development to the two-cell stage was dramatically decreased (Fig. 5 and Table 2). In oocytes electroporated with siControl, there was no decrease in the maturation, fertilization, or developmental rate compared to oocytes that did not undergo electroporation.

**Fig. 5.**
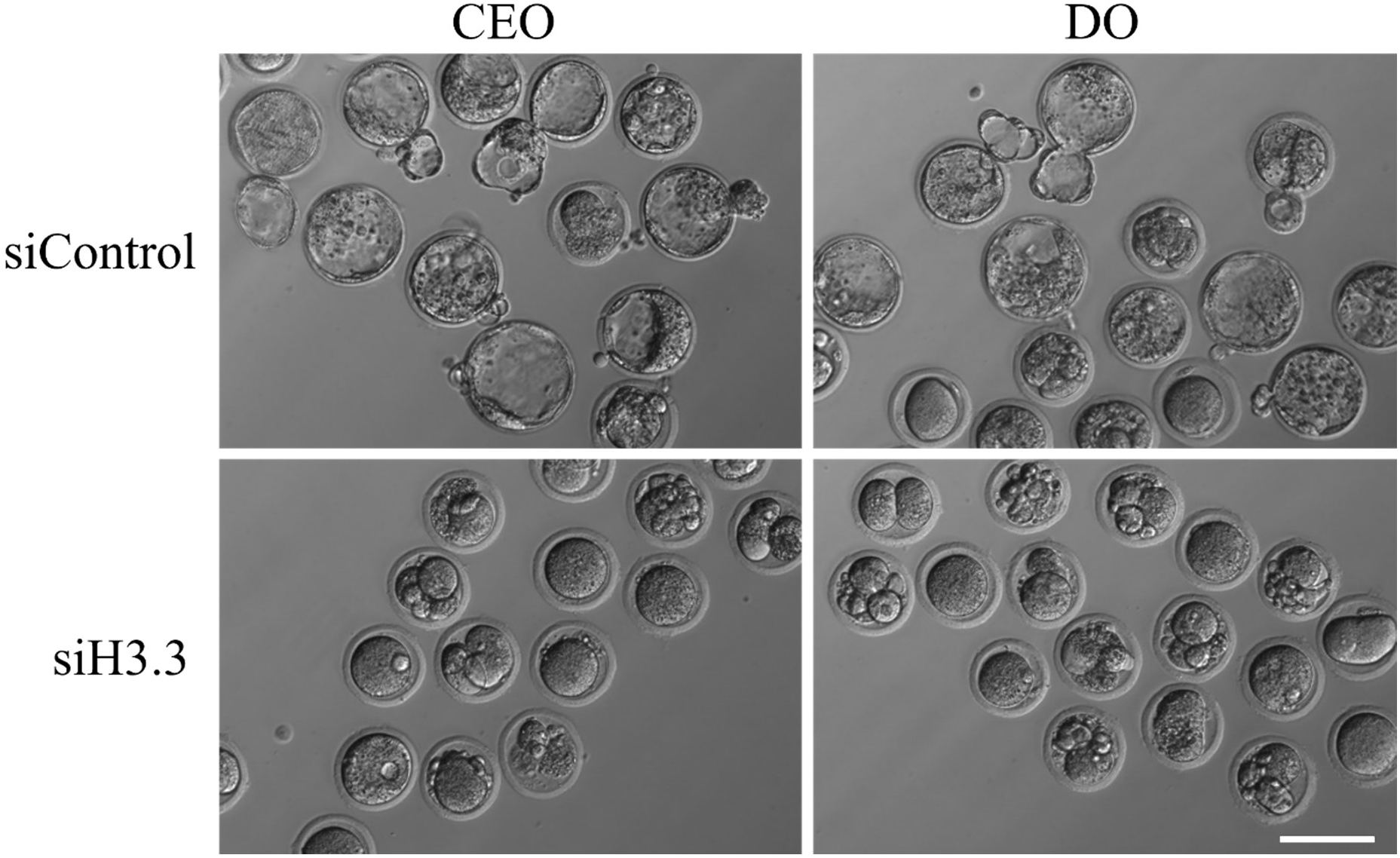
Developmental competence of embryos electroporated with siH3.3 at the germinal vesicle (GV) stage. Representative photos are shown of embryos electroporated with either *siH3.3* or siControl at the GV stage. Embryos were photographed at 96 hpi. CEO: cumulus-enclosed oocytes. DO: denuded oocytes. Scale bar, 100 μm.

**Table 2.** Effects of electroporation of siH3.3 on the developmental competence of CEOs and DOs.

| Treatment of<br>GV oocytes | No. of<br>GV-stage oocytes | No. (%) of<br>MII-stage oocytes | No. (%) of embryos<br>with two pronuclei | No. (%) of<br>2-cell embryos | No. (%) of<br>4-cell embryos | No. (%) of<br>morulae | No. (%) of<br>blastocysts | % of blastocysts per<br>fertilized embryo |
| --- | --- | --- | --- | --- | --- | --- | --- | --- |
| Un-electroporated CEO | 60 | 58 (96.7)<sup>a</sup> | 31 (51.7)<sup>a</sup> | 27 (45.0)<sup>a</sup> | 24 (40.0)<sup>a</sup> | 25 (41.7)<sup>a</sup> | 24 (40.0)<sup>a</sup> | 77.4<sup>a</sup> |
| siControl CEO | 64 | 64 (100.0)<sup>a</sup> | 23 (35.9)<sup>a</sup> | 23 (35.9)<sup>a</sup> | 20 (31.3)<sup>a</sup> | 18 (28.1)<sup>a</sup> | 15 (23.4)<sup>a</sup> | 65.2<sup>a</sup> |
| siH3.3 CEO | 64 | 64 (100.0)<sup>a</sup> | 24 (37.5)<sup>a</sup> | 4 (6.2)<sup>b</sup> | 1 (1.5)<sup>b</sup> | 0 (0.0)<sup>b</sup> | 0 (0.0)<sup>b</sup> | 0.0<sup>b</sup> |
| Un-electroporated DO | 60 | 57 (95.0)<sup>a</sup> | 24 (40.0)<sup>a</sup> | 23 (38.3)<sup>a</sup> | 18 (30.0)<sup>a</sup> | 16 (26.7)<sup>a</sup> | 13 (21.7)<sup>a</sup> | 54.2<sup>a</sup> |
| siControl DO | 63 | 61 (96.8)<sup>a</sup> | 26 (41.3)<sup>a</sup> | 23 (36.5)<sup>a</sup> | 23 (36.5)<sup>a</sup> | 20 (31.7)<sup>a</sup> | 16 (25.4)<sup>a</sup> | 61.5<sup>a</sup> |
| siH3.3 DO | 55 | 54 (98.2)<sup>a</sup> | 28 (50.9)<sup>a</sup> | 7 (12.7)<sup>b</sup> | 1 (1.8)<sup>b</sup> | 0 (0.0)<sup>b</sup> | 0 (0.0)<sup>b</sup> | 0.0<sup>b</sup> |

Different letters (a and b) in the same column represent significant differences ( $P$ value < 0.05).

## Discussion

This study demonstrated reduced developmental competence of DOs after fertilization, which is consistent with previous reports (Shioya et al., 1988; Liu and Aoki, 2002; Kikuchi et al., 2016). The proportion of oocytes with non-surrounded nuclear (NSN)-type chromatin configurations was shown to be higher in DOs than in CEOs (De La Fuente and Eppig, 2001; Liu and Aoki, 2002). NSN-type oocytes are often unable to complete meiosis, and the reduced developmental competence of DOs may be due to the predominance of these oocytes (Debey et al., 1993; Zuccotti et al., 2002). DOs are thought to be generated by apoptosis of granulosa and cumulus cells during folliculogenesis, a phenomenon called atresia (Osman, 1985; Gougeon and Testart, 1986; Hirshfield AN, 1991). This process may disrupt the transzonal cytoplasmic processes between oocytes and cumulus cells, thus reducing the supply of factors such as cAMP that control oocyte maturation and causing a decrease in DO quality (Sutton et al., 2003). Our results showed that dCEOs had dramatically lower fertilization competence than CEOs. The fertilization and developmental rates of dCEOs vary between reports (Schroeder and Eppig, 1984; Chang et al., 2005; Miki et al., 2006; Mahmodi et al., 2009; Zhou et al., 2016), which may be due to differences between mouse strains, culture media, and extent of cumulus cell removal. Our study showed that the addition of cumulus cells to dCEOs partially restored the fertilization and developmental rates, supporting the importance of oocyte-cumulus cell communication (Sutton et al., 2003) and suggesting that dCEOs can be effectively used in this method. These results indicate that CEOs more accurately mimic in vivo conditions than DOs or dCEOs when studying post-fertilization developmental rates.

We found that electroporation could be used to introduce siRNA into CEOs and DOs and thereby suppress target genes. However, in the case of electroporation of CEOs, siRNAs were also introduced into cumulus cells; thus, caution should be taken when targeting genes that are expressed in cumulus cells. Although the incorporation rate was higher with longer pulse lengths, the knockdown efficacy was independent of pulse length, suggesting that a sufficient amount of siRNA can be introduced for gene knockdown in mouse preimplantation embryos regardless of pulse length. Only one study examined the mechanisms of embryo development using antisense oligonucleotide injection at the GV stage followed by in vitro fertilization (Arand *et al*., 2022). This may be because DOs and dCEOs with extremely low fertilization and developmental competence are used; therefore, it is difficult to secure enough fertilized eggs for analysis. The electroporation method allowed us to introduce siH3.3 into DO and CEO and to evaluate the impact of H3.3 on developmental competence after fertilization. It has been reported that H3.3 is preferentially incorporated into the paternal pronucleus and that knockdown of H3.3 in GV-stage oocytes causes abnormal pronuclear formation (Inoue and Zhang, 2014). Our results showed that H3.3 was already present in GV-stage oocytes, as previously reported (Akiyama *et al*., 2011; Nashun *et al*., 2015), and H3.3 localization in the maternal pronucleus was unchanged by the introduction of siH3.3, suggesting that H3.3 incorporated into the maternal genome is tightly bound to the genome even during the transition from GV to MII. In addition, siH3.3 electroporation into GV-stage oocytes inhibited H3.3 incorporation into the paternal pronucleus after fertilization, indicating that maternal H3.3 protein is repeatedly translated and degraded in the oocyte and that the depletion of H3.3 mRNA by siH3.3 causes a gradual decrease in H3.3 protein during oocyte maturation. Electroporating GV-stage oocytes with siH3.3 also reduced the development competence after fertilization, showing that the incorporation of H3.3 into the paternal pronucleus is required for subsequent embryonic development.

The method of electroporating siRNA into GV-stage oocytes described in this study is a powerful tool to analyze the function of maternal factors, and it allowed us to examine the involvement of these factors in embryonic development after fertilization. This method may also be applicable to genome editing involving the introduction of guide RNA and Cas9 protein, which may allow us to generate maternal allele–specific knockout mice. In summary, we comprehensively analyzed the fertility and developmental competence of CEOs, DOs, and dCEOs, and developed an electroporation method for introducing siRNA into CEOs to genetically analyze fully grown oocytes in a manner that does not compromise their developmental potential.

## Author contribution statement

TY, SH, SI, and NM conceived the study. TY, SH, II, and MS performed the experiments. TY analyzed the data. TY, SH, SI, and NM wrote the manuscript, and all authors reviewed the manuscript.

## Acknowledgments

The authors thank all the members of the team for providing technical support and valuable suggestions.

